# BacTaxID: A universal framework for standardized bacterial typing

**DOI:** 10.64898/2025.12.09.693184

**Authors:** Miguel D. Fernández-de-Bobadilla, Val F. Lanza

## Abstract

Bacterial strain typing is key to surveillance, outbreak investigation and microbial ecology, yet current systems remain species-specific, reference-dependent and lack a universal, interpretable metric of genomic relatedness. Here, we introduce BacTaxID, a fully configurable, whole-genome k-mer-based framework that encodes each genome as a numeric sketch and organizes strains into hierarchical clusters with user-defined similarity thresholds. BacTaxID distances are strictly proportional to Average Nucleotide Identity (ANI), providing a direct quantitative link between vectorial typing and genome-wide divergence. Applied to 2.3 million genomes from “All the Bacteria” database across 67 genera, BacTaxID demonstrates universal concordance species and sub-species classification systems, while capturing finer strain-level diversity than traditional reference-based approaches. In simulated surveillance and real outbreak datasets, BacTaxID reproduces SNP and cgMLST-based definitions while enabling rapid, scalable screening. Precomputed genus-level schemes and an open implementation provide a practical, genus-agnostic alternative to classical typing systems for standardized bacterial classification.

## Introduction

Bacterial typing has undergone a paradigm shift from traditional phenotypic methods to whole-genome sequencing (WGS), becoming indispensable for modern epidemiology, outbreak detection, and surveillance ^1–3^. Current gold standards rely heavily on Multilocus Sequence Typing (MLST) and its high-resolution extensions, core-genome MLST (cgMLST) and whole-genome MLST (wgMLST). While MLST utilizes 6-9 housekeeping genes to define portable Sequence Types (STs), cgMLST and wgMLST expand this concept to thousands of loci, offering significantly finer discrimination^2–5^. Decentralized approaches like dcgMLST^6^ employ hash-based allele identifiers to eliminate reliance on curated nomenclature, though they still necessitate shared reference datasets for meaningful comparison. However, fundamental limitations persist: these schemes remain inherently species-specific, creating silos of non-interoperable data^4,7,8^. Furthermore, standard MLST identifiers are arbitrary and phylogenetically opaque, closely related strains can bear unrelated ST numbers due to minor allelic variations^4,9^. Additionally, cgMLST distances tend to saturate rapidly, restricting their utility for broader taxonomic or inter-species comparisons where SNP-based metrics retain discriminatory power ^10,11^.

A critical challenge with high-resolution methods like cgMLST and wgMLST is that they often produce hyper-specific labels that hinder effective data exchange between researchers, laboratories, and public health organizations. Hierarchical frameworks such as HierCC^12^ and LIN^13,14^ have attempted to resolve these issues by organizing strains into nested similarity levels based on cgMLST distances, aiming to group samples into more stable and communicable clusters. While promising, these systems frequently generate unwieldy, long numerical nomenclatures that still impede effective communication compared to simple ST labels. Critically, many rely on single-linkage clustering, a method vulnerable to “chaining artifacts”^15^, where intermediate genomes artificially bridge phylogenetically distinct lineages, obscuring true epidemiological structure. The immense diversity in mutation rates and recombination frequencies across bacterial taxa further complicates the definition of universal thresholds, resulting in a fragmented landscape of isolated typing systems that require intensive, species-by-species validation.

To address these challenges, we introduce BacTaxID, a universal, genome-based typing framework that unifies scalable classification with biologically meaningful nomenclature. By utilizing whole-genome k-mer profiling combined with a novel pseudo-clique-based hierarchical clustering algorithm, BacTaxID prevents the merging of distant lineages and eliminates chaining artifacts common in single-linkage methods. This reference-free approach derives genomic signatures directly from sequence content, providing a single, consistent metric of relatedness applicable across all bacterial species. BacTaxID generates compact, interpretable identifiers that retain the simplicity of MLST while transparently encoding hierarchical relationships, enabling immediate assessment of strain relatedness without external database lookups. Validated on 2.3 million genomes from the *All the Bacteria* database^16^ spanning 67 genera, the system demonstrates high performance and scalability for global surveillance. Pre-computed schemes and tools available at www.bactaxid.org offer a practical, genus-agnostic solution that bridges the gap between automated classification and standardized typing for clinical and evolutionary applications.

## Results

### Hierarchical Typing Algorithm

BacTaxID implements a graph-theoretic hierarchical clustering approach for automated bacterial genome typing, integrating sketch-based distance estimation with iterative level-by-level assignment to generate scalable nomenclature. The system employs Binwise Densified MinHash^17,18^ combined with ntHash^19,20^ for efficient k-mer hashing, decomposing genomes into compact sketch representations without reference bias. Pairwise distances are estimated via Jaccard similarity and transformed to ANI following the equation implemented in MASH^21^. This sketch-based method achieves computational efficiency orders of magnitude higher than alignment-dependent methods while maintaining a strong correlation with evolutionary distances. By restricting comparisons at subsequent levels (L_0_ to L_x_) to classifiers sharing identical typing codes, the hierarchical search strategy reduces computational complexity from O(N) to O(log(N)) or O(N^1/2^), enabling tractable processing of expanding databases.

Conceptually, BacTaxID’s representation parallels cgMLST: both encode genomic information as fixed-length numeric vectors. However, unlike allele-based systems, BacTaxID’s vector elements represent reference-free k-mer hash bins. This eliminates dependence on predefined locus databases and ensures that pairwise vector comparisons directly correlate with genome-wide evolutionary distances measured by Average Nucleotide Identity (ANI), providing a quantitative phylogenetic relationship.

At each hierarchical level, the algorithm compares the query genome against “classifier” references. If the distance falls below the level-specific threshold, the query inherits the reference typing code. The algorithm then checks if the query matches a sufficient proportion of cluster members (defined by *click_threshold*). If this condition is met and the cluster size is below the maximum capacity (*reference_size*), the query becomes a “*classifier*” genome, serving as a reference for future assignments. If the cluster is full, the query is designated as a “*satellite*”. This distinction prevents outliers or hypermutants from acting as references, maintaining robust cluster definitions while allowing valid assignment to saturated groups.

When no existing cluster matches, the system initiates de novo cluster formation using graph-theoretic clique detection. It constructs a distance graph of unclassified genomes sharing the same parent code and identifies maximal cliques: complete subgraphs where all members satisfy pairwise distance thresholds. Valid cliques meeting minimum size requirements establish novel clusters, assigning a new hierarchical code to all members. This approach ensures high internal cohesion and prevents chaining artifacts common in single-linkage clustering. Strict monotonicity constraints ensure that codes at finer levels (L_i+1_) consistently refine groupings from coarser levels (L_i_).

BacTaxID outputs a unified DuckDB database file consolidating typing information, scheme parameters, and the complete k-mer sketch database. This self-contained, portable format supports reproducibility and interoperability via DuckDB’s API ecosystem (SQL, Python, R etc…). The architecture allows both static classification against established schemes and dynamic updates through continuous clustering. Key parameters, including hierarchical levels and similarity thresholds, are user-configurable, allowing adaptation to user needs.

#### All The Bacteria dataset

To demonstrate BacTaxID’s capabilities, we analyzed the All the Bacteria (ATB) database ^16^, a large-scale, open-access project that assembles and catalogs millions of bacterial and archaeal genomes from public repositories. The ATB collection comprised 2.4 million quality-controlled genomes as of August 2024, providing a standardized resource for comparative genomic analysis. We selected all genera containing more than 500 genomes, yielding a comprehensive dataset of 2.3 million genomes distributed across 67 genera and 3,926 species.

Individual typing schemes were generated for each genus using identical parameterization: six hierarchical clustering levels based on ANI thresholds of 96%, 98%, 99%, 99.5%, 99.9%, and 99.99%. Consequently, each genome is assigned a hierarchical typing code composed of up to six integers separated by periods (e.g., 1.3.1.8.12.1), where each integer represents the cluster assignment at a successive level of resolution. All schemes are publicly available at https://zenodo.org/records/17791772, and comprehensive information including the complete dataset, metadata from All the Bacteria, classification results, and interactive exploration tools are freely available at www.bactaxid.org. This unified platform provides researchers with seamless access to all BacTaxID classification data, enabling integration with external analyses and supporting both surveillance applications and evolutionary comparative studies.

We selected the genus as the operational taxonomic level for scheme construction due to ongoing controversies surrounding bacterial species delimitation criteria ^3,22,23^. By initiating classification at the genus level, we allow BacTaxID’s hierarchical framework to resolve infrageneric diversity without imposing a priori species boundaries. Additionally, BacTaxID, like other k-mer-based distance estimation methods, exhibits reduced accuracy at genomic similarities below approximately 85% ANI. The relationship between ANI and Jaccard index follows the transformation ANI = 1-1/k* log (2 J /(1 + J)) ^21^, where k represents k-mer size and J denotes the Jaccard index. As shown in Supplementary Figure 1, this relationship maintains approximate linearity at Jaccard values exceeding 0.1, corresponding to ANI > ∼85%, but deviates substantially at lower similarities. This constraint necessitates establishing the genus as the minimum taxonomic level for reliable BacTaxID classification, as intergeneric comparisons frequently fall below the method’s effective resolution threshold.

Theoretically, the BacTaxID design improves classification performance as the dataset grows. The *click_size* parameter, which requires a minimum number of members to establish a new cluster, is intended to increase the completeness of the typing scheme by preventing the formation of spurious, low-density clusters in larger datasets. Here, we define completeness as the proportion of samples assigned a definitive code at each hierarchical level. Across the 2.3 million genomes analyzed, completeness exhibited high variability between levels, ranging from near-universal assignment (>98%) at the broader L_0_ (96% ANI) resolution to significantly lower rates at the finest L_5_ (99.99% ANI) level (Figure 1B). Completeness shows weak correlation with the number of sequences (Figure 1C), strengthening at L_3_ and L_4_ levels

**Figure 1.**
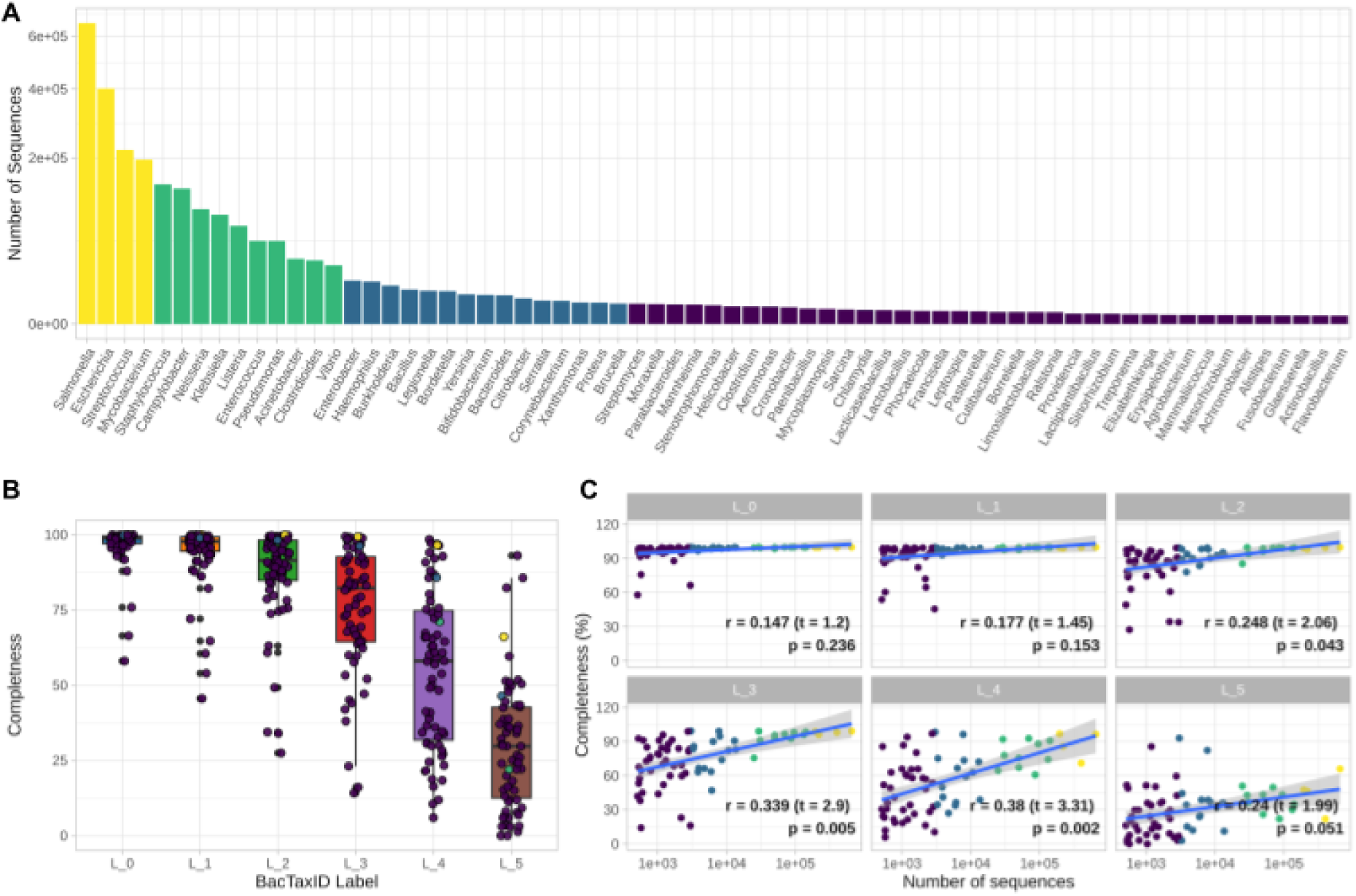
Dataset composition and typing scheme performance across bacterial genera. (A) Distribution of 2.3 million genomes across 67 genera from the All the Bacteria database, ordered by sample size. (B) Completeness distribution at each BacTaxID hierarchical level (L_0_ through L_5_, corresponding to ANI thresholds from 96% to 99.99%). Completeness indicates the percentage of genomes assigned to clusters at each resolution level. (C) Correlation between typing scheme completeness and genus-level dataset size across all six hierarchical levels. Each point represents a single genus, with colors denoting BacTaxID levels. Regression lines demonstrate positive correlation between sample size and scheme completeness, particularly at finer resolution levels. Pearson correlation coefficients (r), t-statistics (t), and p-values are displayed for each level.

### Multilevel Resolution of Bacterial Population Structure

To validate the biological relevance of our hierarchical scheme and assess concordance with established frameworks, we evaluated BacTaxID against consensus classifications in *Escherichia* and *Salmonella*, the two most abundant genera in All the Bacteria (Figure 2). We computed Normalized Mutual Information (NMI) ^24^ between each hierarchical level and reference schemes to identify optimal alignment. Strong concordance emerged between species designations in the ATB database and L> clusters (96% ANI), reflecting the established 95% ANI species threshold^25^. However, some genomes classified as *Escherichia coli* and *Salmonella enterica*, the dominant species in each genus, clustered distinctly within different L_0_ groups (Figure 2D, 2F), suggesting previously unrecognized or inadequately characterized pseudo-species not yet formally described.

**Figure 2.**
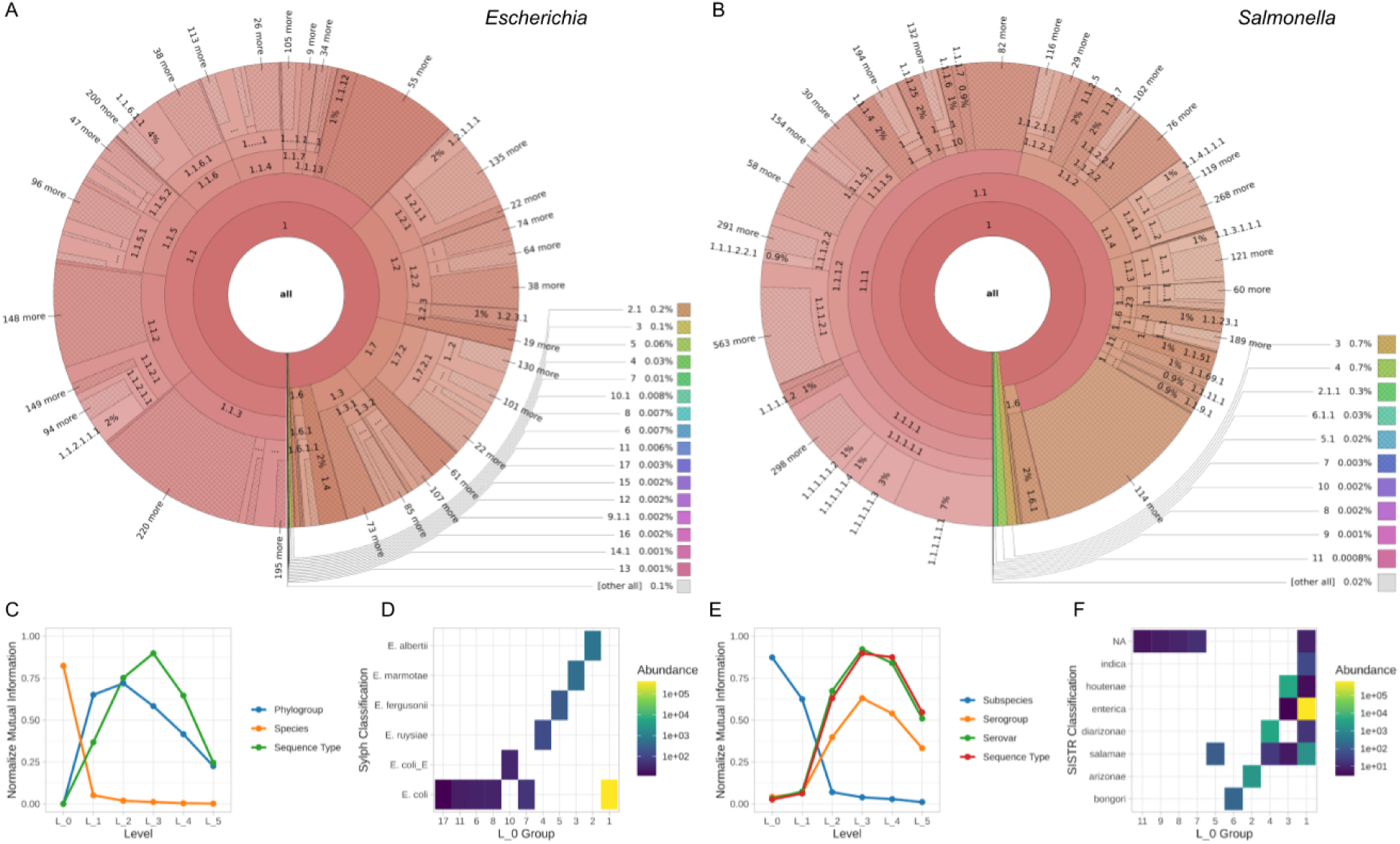
Population structure and agreement with established typing schemes in *Escherichia* and *Salmonella.* (A,B) Krona plots depicting the hierarchical population structure of *Escherichia* and *Salmonella* genera, respectively, as resolved by the BacTaxID typing scheme. Each concentric ring represents a successive hierarchical level (L_0 through L_5), with sector angles proportional to the relative abundance of genomes assigned to each cluster. Numerical labels indicate cluster identifiers, with annotations showing the number of additional clusters collapsed for visualization clarity. (C,E) Normalized Mutual Information (NMI) between BacTaxID hierarchical levels and established typing schemes for *Escherichia* and *Salmonella,* respectively. The x-axis denotes BacTaxID levels (L_0 through L_5), while the y-axis represents NMI values ranging from 0 (no agreement) to 1 (perfect concordance). Colored lines represent different reference typing systems: phylogroups, species, and sequence types for *Escherichia;* and subspecies, serovars, and serogroups for *Salmonella.* Higher NMI values indicate stronger correspondence between BacTaxID clusters and established classifications. (D,F) Species distribution within the first hierarchical level (L_0, 96% ANI threshold) for *Escherichia* and *Salmonella,* respectively. Tile plots display the number of genomes per species (y-axis, styled as classification) stratified by L_0 cluster assignment (x-axis). Color gradients represent genome abundance per species, illustrating the degree of species-level heterogeneity captured within coarse-resolution BacTaxID clusters.

For *E. coli*, a species with extensive phylogenetically diversity^26^, well-established phylogroup assignments showed strong correlation with L_2_ clusters (Figure 2D). At L_3_, BacTaxID clusters demonstrated high concordance with MLST assignments, with NMI values indicating robust agreement between both methods (Figure 2C). This reflects that fine-scale differentiation at this resolution aligns with epidemiologically informative MLST typing. Two additional levels (L_4_ and L_5_, corresponding to 99.5% and 99.99% ANI) capture sub-clonal diversity and micro-epidemiological variants beyond MLST.

For *Salmonella*, we compared BacTaxID against the Salmonella In Silico Typing Resource (SISTR) ^27^ which provides subspecies, serogroup, serovar and MLST information. Similar to *Escherichia*, L_3_ level showed strong NMI concordance with serovar and MLST typing (Figure 2E).

To assess the generalizability of our findings beyond *Escherichia* and *Salmonella*, we extended our comparative analysis by adding the 14 most abundant genera in the ATB database (*Acinetobacter, Campylobacter, Clostridioides, Enterobacter, Enterococcus, Haemophilus, Klebsiella, Listeria, Mycobacterium, Neisseria, Pseudomonas, Staphylococcus, Streptococcus* and *Vibrio*). This set comprised 2.2 million genomes, representing approximately 92% of the collection (Figure 3).

**Figure 3.**
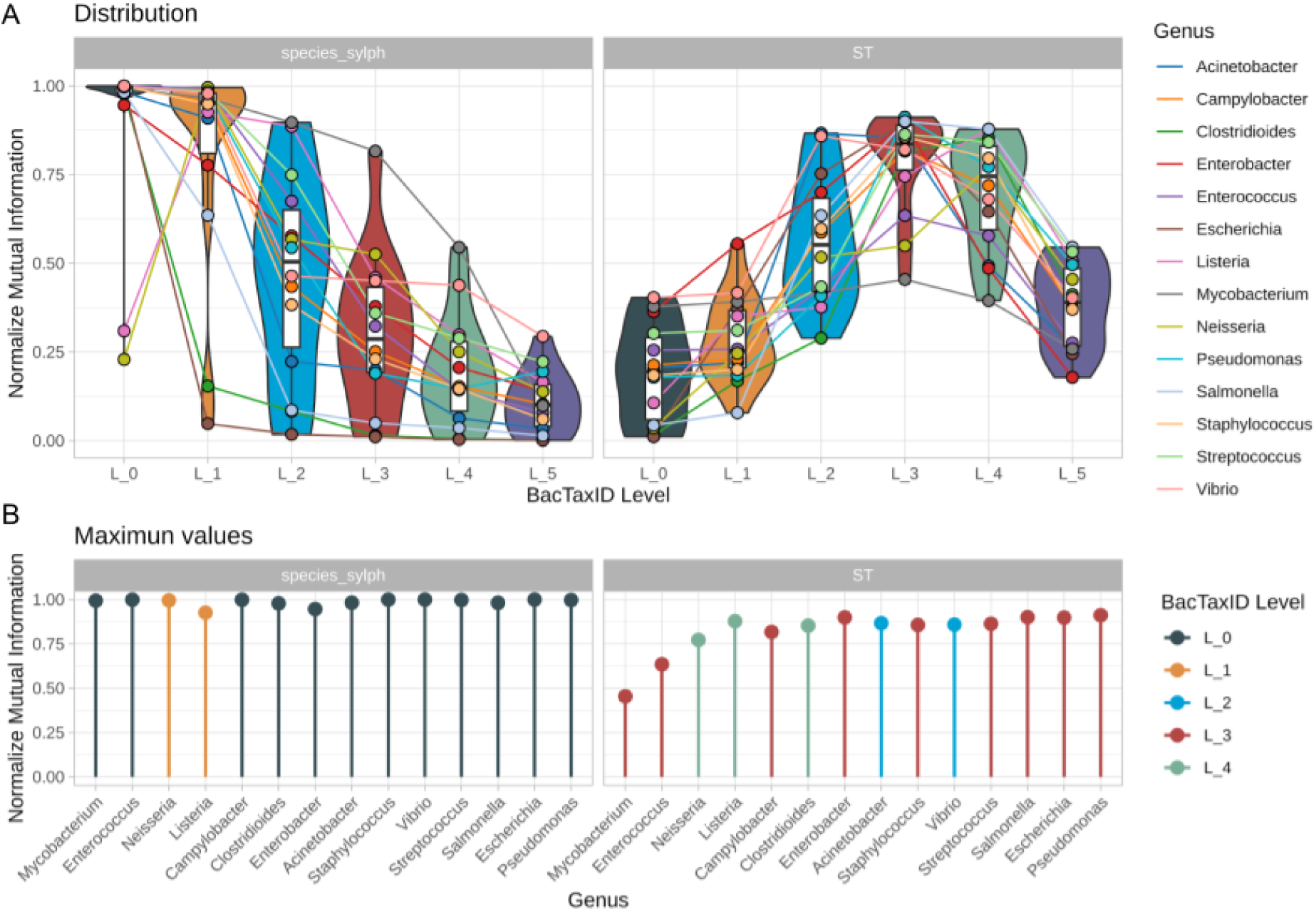
Consistency of BacTaxID typing performance across bacterial genera. (A) Normalized Mutual Information (NMI) between species designations and MLST versus BacTaxID hierarchical levels across 14 most abundant genera of ATB database. Violin plots display the distribution of NMI values per level. (B) Maximum NMI values comparing species and MLST classifications for each genus. Colors correspond to the BacTaxID level achieving the maximum NMI value.

BacTax clusters at L_0_ and L_1_ showed generally high concordance across all evaluated genera, indicating that the coarsest hierarchical levels broadly capture species-level genetic boundaries (Figure 3A). On other hand, MLST assignments showed a consistent peak in concordance at L_3_ (99% ANI threshold) across most genera (Figure 3B), with substantial variation in NMI trajectories at finer resolutions. The observed peak in MLST agreement at L_3_ across phylogenetically diverse bacterial lineages is consistent with this level corresponding to an epidemiologically informative resolution.

In order to evaluate the capabilities of BacTaxID versus the cgMLST we have created cgMLST schemes for the 14 selected genera. As cgMLST operates at the species level rather than the genus level, we selected a homogeneous diversity panel of genomes representing the main BacTaxID group (1.x.x.x.x.x) that correspond to major bacterial species, namely *Escherichia coli*, *Salmonella enterica, Acinetobacter baumannii*, among others. The cgMLST scheme size varied substantially across genera, ranging from 714 genes in *Enterococcus faecium* to 1,291 genes in *Campylobacter jejuni*, reflecting the genomic diversity within each taxon (Figure 4).

**Figure 4.**
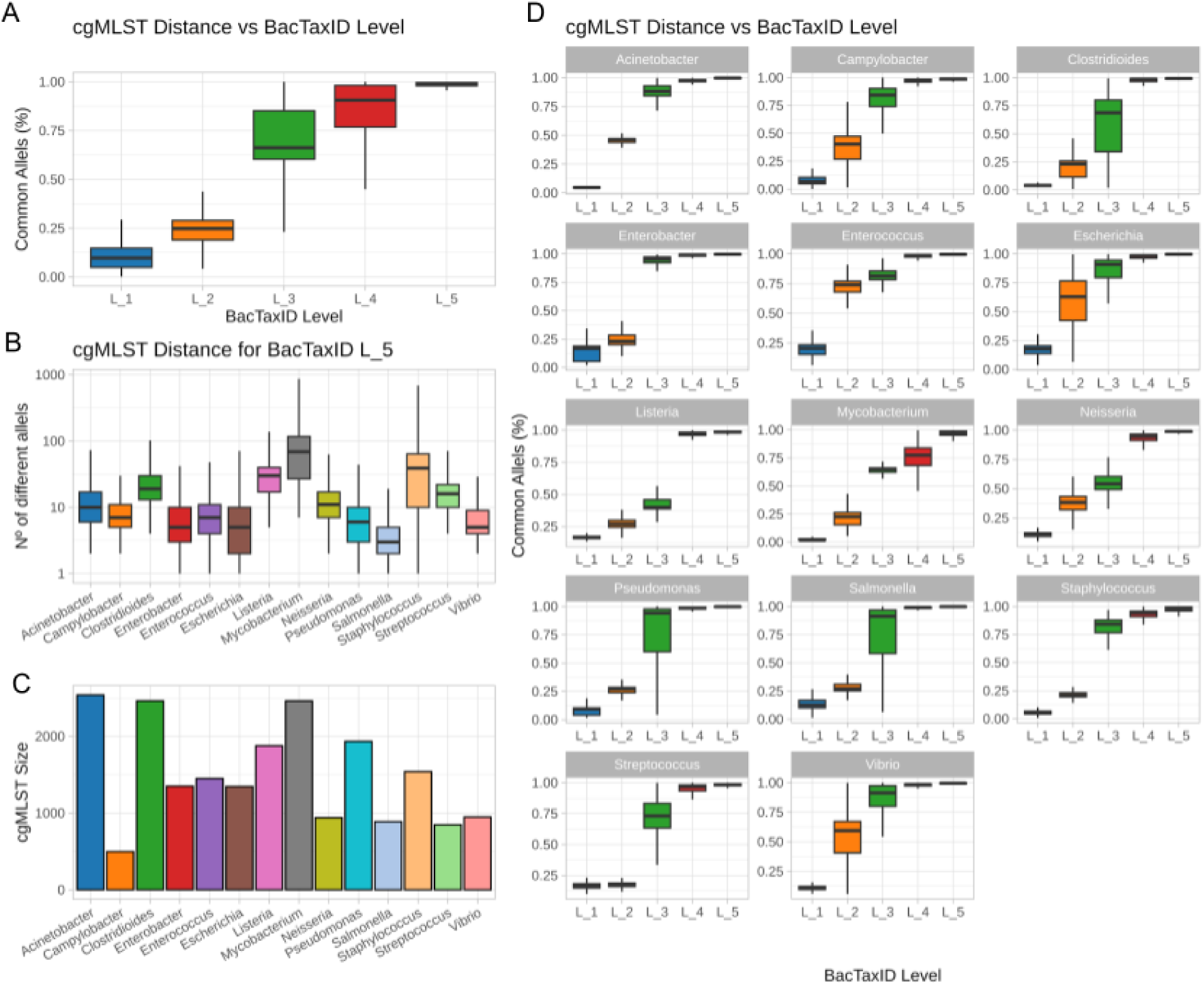
Correlation between BacTaxID taxonomic levels and cgMLST distances across bacterial genera. (A) Percentage of cgMLST common alleles versus BacTaxID level. (B) Number of different alleles between isolate pairs assigned to the same BacTaxID L_5 level, stratified by bacterial genus. (C) cgMLST scheme size for each genus. (D) Percentage of common alleles versus BacTaxID level stratified by genus.

To assess the correlation between BacTaxID taxonomic levels and cgMLST genetic distances, we analyzed the percentage of common alleles across the five BacTaxID hierarchical levels (L_1_ to L_5_) and compared them with cgMLST distance metrics. Our results demonstrated a progressive increase in genetic similarity from L_1_ to L_5_ (Figure 4A), with common alleles ranging from approximately 10% at the most distant level (L_1_) to >95% at the most specific taxonomic level (L_5_). When examined at the species level, individual isolates assigned to the same BacTaxID L_5_ group showed considerable variation in the number of differing alleles depending on the bacterial genus, highlighting the differential resolution provided by BacTaxID across different taxa (Figure 4B and D).

#### Hierarchical Resolution Optimization for Epidemiological Surveillance and Outbreak Investigation

To demonstrate BacTaxID’s utility in real-world biomedical and public health contexts, we evaluated its performance in both surveillance and outbreak detection scenarios.

For the surveillance simulation, we selected 30,000 genomes of *E. coli,* from the ATB database and stratified them by prevalence frequencies from a meta-analysis of extraintestinal pathogenic *E. coli* (ExPEC) infections^28^, enabling direct comparison between MLST-based estimates and BacTaxID’s hierarchical approach (Figure 5A, 5B). At broadest resolution (L_1_), a dominant cluster (1.2) comprised 52% of the population, encompassing ST131, ST12, ST127, ST73 and ST95, followed by clusters 1.1 (22.3%) and 1.3 (17.3%) (Figure 5B). At intermediate resolution (L_2_), cluster 1.2 subdivided into 1.2.1 (25.45%, almost entirely ST131) and 1.2.2 (32.34%, retaining other STs) (Figure 5C). At L_3_, cluster distributions closely matched MLST prevalence (Figure 5D).

**Figure 5.**
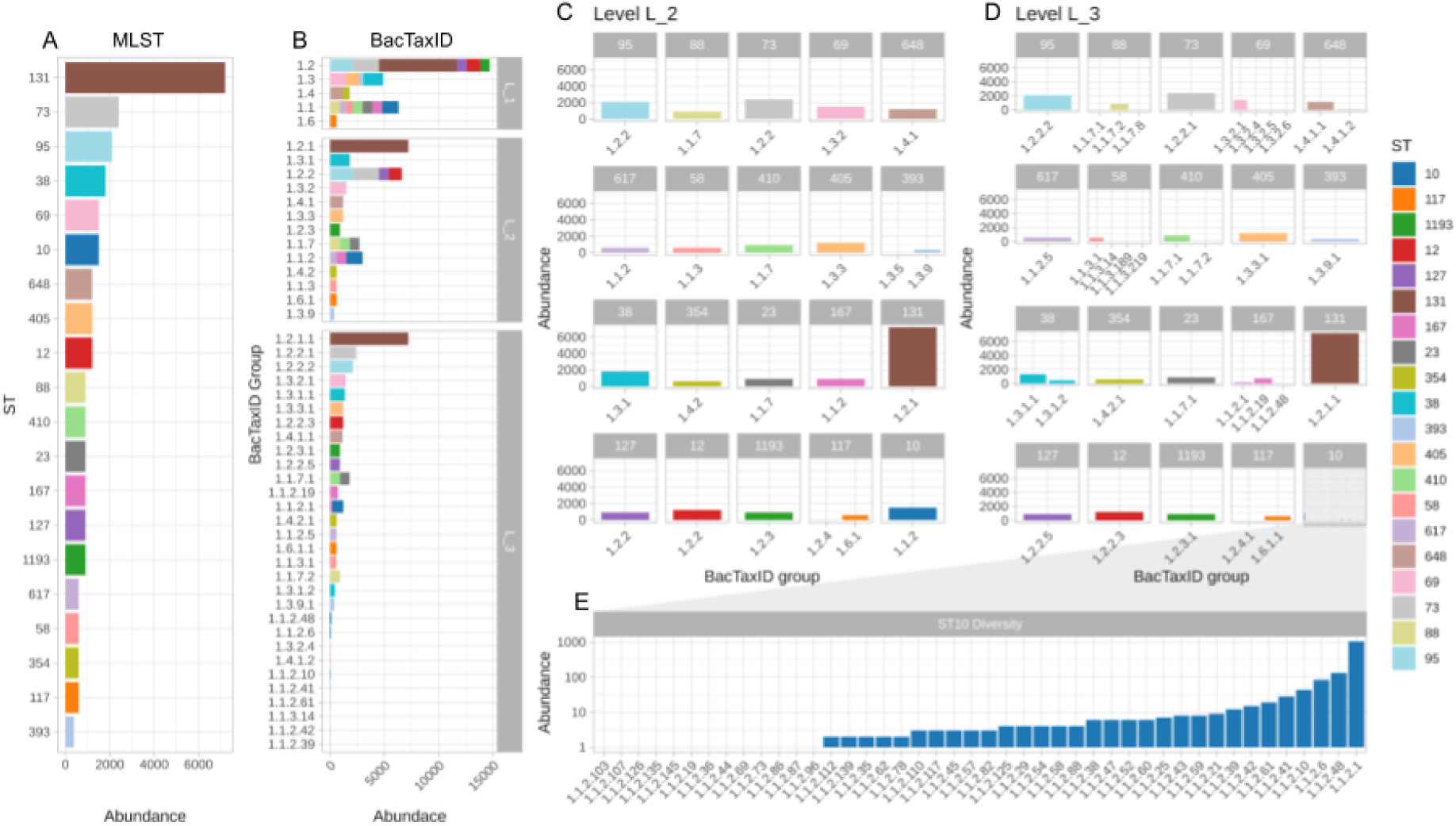
Multi-resolution strain classification in a prevalence survey context. (A) Distribution of MLST sequence types (ST) among *Escherichia* co// isolates from the All the Bacteria database stratified according to reported prevalence frequencies, ranked by abundance and colored by ST designation. (B) BacTaxID hierarchical clustering of the same isolate collection, displaying cluster assignments at levels L_1_, L_2_, and L_3_alongside their relative abundances. Only clusters containing more than 10 members are displayed for visualization clarity, demonstrating alternative resolution perspectives for the same genomic data. (C, D) Detailed distribution of BacTaxID hierarchical levels stratified by individual ST, comparing L_2_ and L_3_ resolutions. Bar plots display the abundance of each BacTaxID cluster nested within every detected ST, illustrating how intermediate hierarchical levels reveal sub-clonal structure not captured by MLST allelic profiling. (E) Fine-resolution structure within ST10, showing the distribution of 39 distinct lower-order BacTaxID groups at higher resolution levels, colored by ST assignment.

Critically, L_3_ analysis revealed individual MLST types are not strictly monophyletic. For example, ST10 segregated across multiple L_3_ clusters (Figure 5E), reflecting substantial pangenomic diversity within a single designation. This polyphyly suggests that MLST-based prevalence estimates may conflate distinct sublineages. By contrast, BacTaxID’s hierarchical resolution allows more nuanced strain characterization, supporting refined risk stratification and epidemiological interpretation.

Outbreak definition relies on a multifactorial assessment incorporating case counts, spatial distribution, temporal clustering, and genomic similarity. In this context, genomic relatedness is typically evaluated using allele differences in cgMLST or Single Nucleotide Polymorphism (SNP) distances, which serve as the gold standards for resolving transmission chains^29–31^. To assess BacTaxID’s capacity to capture this fine-scale genomic variation, we computed the SNP density among isolates assigned to identical BacTaxID clusters across 10 representative clusters per species (Figure 6A). At the L_4_ level (99.5% ANI), the median SNP density ranged from 6 (3-18) SNPs/Mb in *Enterococcus* to 183 (35-344) SNPs/Mb in *Neisseria*. At the finer L_5_ level (99.99% ANI), resolution improved significantly, with median SNP densities dropping to 3 (0-14) SNPs/Mb in *Escherichia* and 29 (14-44) SNPs/Mb in *Mycobacterium*. These results demonstrate that BacTaxID captures high-resolution genomic variation, with L_5_ offering sub-clonal discrimination comparable to SNP-based analysis.

**Figure 6.**
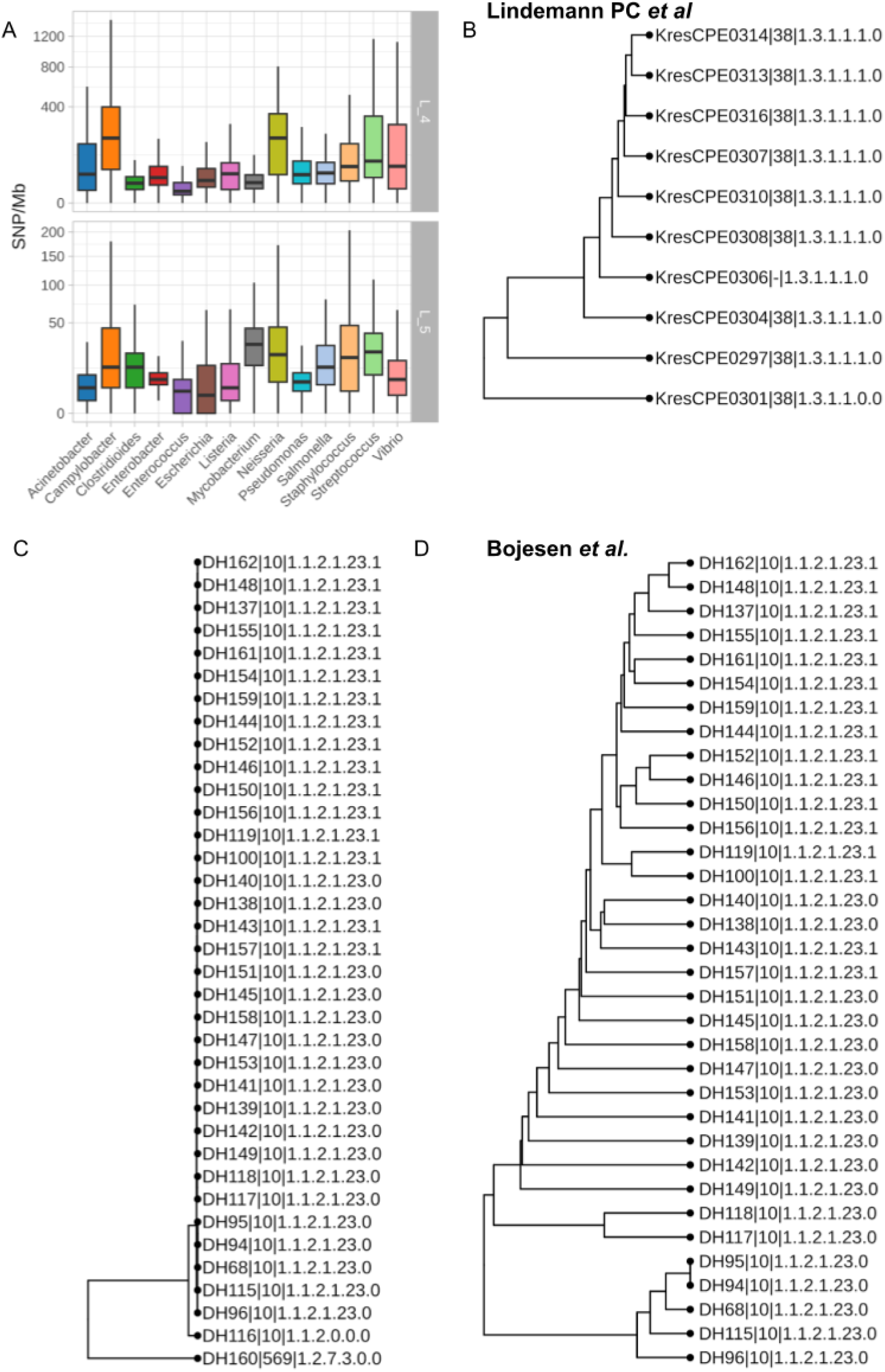
Outbreak detection capability and genomic equivalence of BacTaxID hierarchical levels. (A) Distribution of single nucleotide polymorphism (SNP) density per megabase within individual BacTaxID clusters at hierarchical levels L_4_, (99.5% ANI) and L_5_ (99.99% ANI) across 10 representative genera(B) Similarity tree of the Lindemann *et al. Escherichia coli* ST38 nosocomial outbreak, annotated with BacTaxID hierarchical codes at L_3_. (C) Similarity tree of Bojesen *et al*. broiler outbreak of *Escherichia coli* ST10 poultry outbreak isolates, displaying the hierarchical classification. (D) Expanded similarity tree of the broiler outbreak isolates, illustrating fine-scale sub-clonal differentiation within the dominant outbreak cluster.

To further validate BacTaxID’s capabilities in real-world outbreak scenarios, we reanalyzed two well-characterized cases. Lindemann et al.^32^ documented a nosocomial *E. coli* ST38 (OXA-244) outbreak across three Norwegian hospitals (July-November 2020; 12 cases), where cgMLST analysis showed 0-8 allelic differences within the outbreak cluster versus 15-62 differences from unrelated ST38 isolates. Similarly, Bojesen et al.^33^ described seven broiler unit outbreaks over 18 months (June 2017-November 2018) involving clonal *E. coli* ST10 (O132:H7), with ≤9 SNP differences observed among 36 isolates. BacTaxID clusters demonstrated strong concordance with these published characterizations (Figure 6B-D), confirming its utility for precise outbreak investigation.

## Discussion

The explosion of genomic databases has made traditional classification systems unsustainable, creating an urgent need for scalable, hierarchical frameworks. Current methods, fragmented into species-specific schemes, cannot keep pace with bacterial discovery nor offer true interoperability^34,35^. BacTaxID addresses this problem through a universal approach based on whole-genome k-mer profiling. By eliminating dependence on predefined loci, our system avoids constant scheme redesign and produces standardized, comparable outputs across clinical, evolutionary, and environmental domains. This hierarchical architecture not only manages massive data volume but resolves species definition controversies by adopting the genus as the operational unit, allowing intrinsic population structure to define biological boundaries naturally.

The current standard, cgMLST, presents critical structural limitations^5,36,37^. First, by relying exclusively on the core-genome, it ignores significant genomic diversity (accessory genes, pathogenicity islands) essential for strain characterization. Second, it is inherently subjective: its resolution depends on the reference strain collection used for scheme construction, biasing results toward well-characterized lineages. Third, our data (Figure 4) demonstrate that cgMLST loses discriminatory power drastically when Average Nucleotide Identity (ANI) falls below 99% (L_2_-L_3_). At this range, loci become invariant or saturate their divergence, compressing distances and obscuring true evolutionary structure.

Prior systems like LinBASE^38^ or HierCC^12^ attempted to hierarchize cgMLST but inherited its fundamental limitations, additionally introducing the chaining problem from single-linkage clustering, where intermediate strains artificially merge distant groups^39,40^. BacTaxID overcomes this through a pseudo-clique-based algorithm guaranteeing internal cohesion, and by operating on the complete genome, maintains strict linear correlation with ANI. This creates a direct quantitative link between vectorial distance and actual genomic divergence, absent in traditional typing schemes.

Our results confirm that BacTaxID provides the universal, standardized framework that microbiology lacked. Rather than replacing cgMLST, BacTaxID functions as a complementary tool that enables hierarchical epidemiological workflows. BacTaxID’s primary strength lies in contextual flexibility: configurable hierarchical levels (L_2_-L_3_ for surveillance, L_4_-L_5_ for outbreak investigation) adapt to diverse epidemiological questions without redesigning classification schemes. In contrast, cgMLST prioritizes precision within species through fixed core-genome loci and established allele cutoffs (typically 0-10 differences) ^41^, providing exceptional discriminatory power for transmission chain identification^5,42^. BacTaxID maintains resolution at high-ANI ranges (99.5-99.99%) where cgMLST saturates, while the hierarchical structure naturally guides cgMLST toward relevant comparisons, creating a stratified workflow where BacTaxID defines global genomic neighborhoods and cgMLST refines local transmission chains within them.

This complementary framework is practical and scalable. BacTaxID serves as the initial screening tool, rapidly partitioning genomes across species and genera with computational efficiency and globally standardized nomenclature, while its portable DuckDB format enables fully decentralized laboratory analysis. Once potential outbreaks are identified via BacTaxID’s hierarchical resolution, targeted cgMLST analysis focuses on defined populations for precision allelic differentiation, and when epidemiological urgency demands forensic-level discrimination, SNP-level phylogenetic analysis can be selectively applied to refine transmission chains. This two-stage approach combines BacTaxID’s universality and speed with cgMLST’s epidemiological interpretability, strengthening outbreak detection and cross-species surveillance while remaining feasible for resource-limited settings.

Our retrospective outbreak analyses demonstrate how BacTaxID complements rather than replaces established methods. In nosocomial E. coli ST38 ^32^ and zoonotic E. coli ST10 ^33^ outbreaks, BacTaxID cluster assignments agreed with cgMLST or SNP-based definitions. Level L_5_ offers sub-clonal resolution comparable to SNP investigation (5.92-27 SNPs/Mb), while the hierarchical design enables rapid prospective surveillance at L_3_-L_4_, providing early warning and guidance for escalation to intensive molecular investigations.

BacTaxID balances practical decentralization with essential standardization through a semi-centralized architecture: portable DuckDB files enable fully autonomous, server-independent genome profiling by individual laboratories, while centralized nomenclature coordination through www.bactaxid.org ensures globally consistent strain nomenclature across surveillance networks, a critical requirement that distributed systems cannot achieve independently. This platform consolidates hierarchical nomenclature, scheme distribution, and standardized outputs for 67 bacterial genera, providing pre-computed schemes integrating species designations and antimicrobial, biocide and metal resistance genes and virulence factor profiling via AMRfinderPlus^43^(data available at All the Bacteria database). Laboratories maintain complete analytical autonomy, performing genome profiling locally at their computational capacity without remote server dependence, while standardized nomenclature ensures nomenclatural stability and enables cross-institutional communication. This architecture facilitates dynamic scheme evolution: as databases expand, updated schemes released through www.bactaxid.org seamlessly accommodate new data without reorganizing existing classifications. With comprehensive schemes covering >2.3 million sequenced bacteria and scalability to millions of genomes, BacTaxID positions itself as the practical tool unifying epidemiological surveillance, outbreak investigation, and microbial ecology under a single standardized yet locally autonomous genomic language.

## Online Methods

### Sketch Generation and Distance Estimation

BacTaxID is implemented in Rust, a systems programming language that provides substantial computational advantages for large-scale bioinformatic workflows. Rust’s memory safety guarantees eliminate entire classes of runtime errors including buffer overflows, use-after-free conditions, and data races, ensuring robust handling of diverse genomic datasets without sacrificing performance. Rust’s native parallelization capabilities through packages like rayon^44^ allow seamless multi-threaded processing of large genome collections, enabling efficient utilization of modern multi-core processors. The resulting BacTaxID implementation achieves high throughput while maintaining strict resource management, making it suitable for processing millions of genomes.

BacTaxID employs a custom implementation combining binwise densified MinHash ^17,18^ with NtHash ^19,20^ for efficient k-mer hashing and genome sketching. The process consists of three integrated stages: genome parsing and preprocessing, k-mer extraction and hashing, and binwise signature construction.

Genome sequences are parsed from FASTA files (or compressed FASTA files) using the *needletail* ^45^ library, a high-performance Rust bioinformatics library that processes sequence records sequentially without loading entire files into memory. For each genome record, sequence normalization is applied to convert sequences to a canonical form (typically converting to uppercase and handling degenerate bases). The genome identifier (filename or sequence header) is extracted and normalized for integration into the hierarchical classification database. This stage establishes the foundation for consistent, reproducible sketch generation across diverse data sources.

Following normalization, each genome undergoes k-mer decomposition via a rolling-hash iterator implemented through the NtHash algorithm. Rather than explicitly generating and storing all k-mers (which would require N - k + 1 memory allocations per genome, where N is genome length), NtHash computes canonical k-mer hashes on-the-fly using a rolling window: as the window advances by one nucleotide, the hash is updated incrementally in constant time by removing the contribution of the departing nucleotide and incorporating the new incoming nucleotide. This streaming approach avoids materializing all k-mers, enabling memory-efficient processing even for large genomes (e.g., several Mb). The k-mer size parameter (denoted k) is configurable and defines the granularity of genomic information captured. Rust’s ownership system and compile-time borrow checking guarantee memory safety throughout this iterative processing, preventing common programming errors while the compiler automatically optimizes the generated machine code for the target hardware. The resulting hash values are 64-bit unsigned integers uniformly distributed across the hash space [0, 2^64^ - 1].

The binwise MinHash algorithm partitions the 64-bit hash space into a predefined number of bins *(sketch_size,* denoted S), where each bin spans approximately T2^64^ / SI contiguous hash values. As k-mer hashes are generated by the rolling iterator, each hash is assigned to its corresponding bin via integer division:

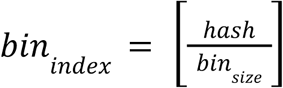

Within each bin, only the minimum hash value encountered during sequence traversal is retained. This direct binning strategy operates in constant time per k-mer (O(1)), avoiding the O(log S) heap operations required by bottom-k MinHash schemes, while maintaining sketch quality comparable to traditional approaches. The resulting sketch is a fixed-length vector of S 64-bit minimum values (one per bin), effectively compressing the entire genome sequence into a compact representation.

The sketch vector is serialized to binary format using the *bincode*^46^ library, which provides fast encoding and low-overhead representation without auxiliary metadata. This serialization enables rapid deserialization during hierarchical clustering operations and facilitates storage in the DuckDB^47^ database. Duplicate k-mers within a sequence are automatically filtered through the streaming process: if the same k-mer appears multiple times, only the first occurrence contributes to the binwise construction, and subsequent duplicates do not affect the bin minimums, this behavior is inherent to the MinHash algorithm and requires no explicit duplicate removal step.

Pairwise genomic distances are computed by comparing sketch vectors: for two sketches S_i_ and S_2_, the number of matching bins (identical minimum hash values) is counted, and the Jaccard similarity is approximated as:

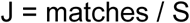

where matches is the count of bins with identical values and S is the sketch size. This Jaccard estimate approximates the intersection-over-union of the underlying k-mer sets, capturing genomic overlap without explicit set operations. The relationship between Jaccard similarity and average nucleotide identity (ANI) is established through the Mash distance formula^27^:

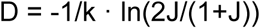

ANI is then calculated as ANI = 1 - D, and clamped to the range [0, 1]. This transformation maintains linearity and accuracy for ANI values ≥85% (corresponding to J ≥ 0.1), which encompasses the vast majority of meaningful bacterial comparisons; below this threshold, approximation error increases substantially, establishing the practical lower bound for BacTaxID’s reliable operation.

Distance computations exploit merge-sort comparison of ordered hash arrays, achieving O(S) complexity per pairwise comparison. Parallelization via *rayon*^48^ enables efficient all-against-all or hierarchical subset comparisons across millions of genomes, reducing wall-clock time to minutes even for massive datasets. In hierarchical searches (described in the Methods section on Hierarchical Clustering), comparisons are restricted to biologically relevant subsets, reducing effective complexity from O(N^2^) to O(N log N) or O(N √N).

All sketch data, including serialized binwise signatures, genome identifiers, and hierarchical classification results, are consolidated into a self-contained DuckDB database file. This unified format eliminates external dependencies on separate sketch files and ensures full reproducibility: any researcher can query results decades later without relying on external services or archived databases. The database supports standard SQL queries as well as interfaces via Python, R, and command-line tools, enabling seamless integration into diverse analytical pipelines. Sketch parameters (k, S) are stored alongside results, ensuring future replicability and enabling cross-validation of findings.

### Hierarchical Clustering and Code Assignment

BacTaxID implements a recursive hierarchical clustering strategy that processes genomes level-by-level, progressively assigning hierarchical classification codes while maintaining biological consistency and computational efficiency. The algorithm operates through two primary pathways: direct assignment to existing clusters (classifier search) or creation of new clusters via graph-theoretic clique detection (“de novo” group formation).

The system processes genomes through a sequence of hierarchical levels (L_0_, L_1_, …, L_N_), each with a predefined distance threshold corresponding to a specific ANI boundary (e.g., L_0_ = 96% ANI, L_3_ = 99% ANI). Thresholds are stored in the database as ordered pairs (*level_name*, *distance_threshold*) and retrieved during initialization. The algorithm traverses levels in ascending order of stringency: coarser resolutions (lower ANI thresholds, earlier levels) are evaluated first, and finer resolutions (higher ANI thresholds, later levels) only when necessary. This order exploits the hierarchical constraint that a genome’s code at level i must be consistent with its code at level i-1: specifically, if a genome is assigned to group 1.3 at L_1_, then its L_0_ code must be 1 (the parent group). This monotonicity constraint is enforced at every assignment step and prevents inconsistent classifications.

Within each cluster at a given level, genomes receive one of two designation states: classifier (C) or satellite (S). Classifier genomes serve as references for subsequent hierarchical searches and new isolate assignments. Satellite genomes are assigned to clusters based on similarity to classifiers but do not themselves serve as references, this distinction prevents spurious clustering caused by outliers, recombinants, or hypermutants that may satisfy distance thresholds but represent atypical lineages. The decision to assign classifier status depends on two parameters: *click_threshold* (typically 80%) and *reference_size* (typically 100 genomes per cluster). Specifically, a candidate genome is promoted to classifier status if: (1) its distance to a reference cluster is below the level threshold, (2) the fraction of reference cluster members within the distance threshold is at least *click_threshold*, and (3) the current cluster size remains below *reference_size*. This criterion prevents indefinite cluster expansion while permitting controlled growth as new genomes with substantial similarity arrive.

When a new query genome is being classified at level i, the algorithm first retrieves all classifier genomes from the database that share the query’s parent-level code (e.g., if the query was assigned to group 1.3 at L_1_, only classifiers with L_1_ code = 1.3 are retrieved). The query sketch is then compared against these reference sketches to compute pairwise distances. If any reference sketch satisfies the level-specific distance threshold, the query is assigned to that cluster, inheriting the cluster’s hierarchical code. The closest reference (best hit) is prioritized, but consistency checks ensure that the reference cluster’s code maintains the required parent-level prefix. If a valid best hit exists and classifier criteria are met, the query is marked as a classifier at that level. This pathway is computationally efficient because the comparison space is restricted to biologically coherent subsets rather than all genomes, achieving O(log N) or O(^N) complexity per level compared to O(N) for flat clustering schemes.

If no valid best hit is identified at a given level (or if the best hit has code = 0 indicating an unassigned reference), the algorithm transitions to graph-theoretic clique detection. In this pathway, the algorithm retrieves all genomes (both classifiers and satellites) that share the query’s parent-level code, computes all pairwise distances between the query and these references, and constructs a distance graph where nodes represent genomes and edges connect pairs whose distance falls below the level threshold. The system then searches for maximal cliques, complete subgraphs where all pairwise distances satisfy the threshold, using iterative deepening and constraint propagation. A clique is considered valid for group formation if its size meets the minimum *click_size* threshold (typically 5 genomes). Upon discovery of a valid clique, the query is assigned a new unique code at that level (the next unused integer for that parent group), and all clique members are updated with this new classification. Importantly, non-clique members in the distance graph receive the same new code but with state = S (satellite), allowing them to be associated with the emerging cluster without influencing future reference searches.

Hierarchical codes are constructed incrementally as the algorithm descends through levels. At L_0_, codes are simple integers (1, 2, 3, …). At subsequent levels, codes are concatenated with dots: if a genome is assigned to group 5 at L_0_, group 3 at L_1_, and group 7 at L_2_, its full hierarchical code is 5.3.7. This hierarchical identifier uniquely specifies the genome’s position in the similarity tree and enables efficient database queries via string prefix matching. The *code_full* field is maintained alongside the numeric code value for both human readability and computational efficiency in hierarchical filtering operations.

The algorithm processes each level sequentially and terminates at a given level under three conditions: (1) a valid classifier-state assignment is achieved and the genome is classified as a classifier, (2) the genome has been assigned via clique detection and new cliques no longer form at finer levels, or (3) no valid distances or cliques are detected at a particular level, leaving that level (and all finer levels) unassigned. This design permits incomplete classifications, where a genome may be fully classified up to L_3_ but lack assignments at L_4_ and beyond. Incompleteness is expected and acceptable when population structure does not support finer-scale clustering, and provides an explicit signal to downstream analyses that higher-resolution typing is not possible or not meaningful for that particular genome within its biological context.

All parameters governing hierarchical classification (number of levels, level-specific thresholds, *click_size*, *click_threshold*, *reference_size*) are user-configurable and stored in the database metadata table upon scheme initialization. This flexibility allows genus-specific or species-specific tuning: organisms with higher evolutionary rates may require coarser thresholds at given ANI boundaries to capture meaningful epidemiological groups, whereas organisms with lower diversity may achieve adequate resolution at finer thresholds. The algorithm scales linearly with the number of query genomes (each processed independently via parallel workflows) and logarithmically with database size per level (due to hierarchical space partitioning), enabling efficient processing of millions of genomes with commodity hardware.

Classification results are stored in a centralized database table that maintains hierarchical information for each genome: the numeric classification code at each level (e.g., 5, 3, 7), the complete hierarchical designation as a concatenated string (e.g., 5.3.7), and the genomic role (classifier or satellite). Database operations employ transactional integrity protocols to ensure that if an error occurs during the writing of classification assignments, the entire operation is rolled back rather than corrupting the database with incomplete or inconsistent data. This prevents cascading errors and maintains data reliability even if processing is interrupted. Additionally, an optional debug mode logs all intermediate computational steps, including genome-to-reference distances and clique detection results, to facilitate retrospective validation of classification decisions and troubleshooting of ambiguous or unexpected assignments.

The update command orchestrates the complete workflow: it loads the SketchManager from the DuckDB database, processes each input FASTA file to generate a query sketch, executes hierarchical classification through iterative level traversal, and persists the resulting sketch and classification codes to the database. This modular design enables efficient incremental updates: new genomes can be added to an existing scheme without re-clustering previously classified samples, maintaining deterministic identifier assignments and enabling long-term stability of nomenclature across laboratory networks.

### Graph construction and clique detection

BacTaxID employs a graph-theoretic framework for de novo cluster formation when a query genome cannot be assigned to an existing group. This process involves dynamic graph construction from pairwise distance data and the application of maximal clique detection algorithms to identify internally consistent groups of genomes.

When direct assignment fails at a given hierarchical level, the algorithm constructs an undirected graph in memory to represent pairwise genomic relationships. First, it retrieves all genomes that share the same parent-level code as the query genome but are not yet classified at the current level (i.e., their hierarchical code for this level is empty). This set includes the query genome itself along with any previously processed genomes that failed to be assigned to a cluster at this resolution. Pairwise distances between all genomes in this unclassified set are computed, and this distance data is inserted as edges into a temporary edges table in the DuckDB database.

The in-memory graph is then built using the *petgraph*^49^ library. Each unique genome identifier is added as a node, and edges are established between nodes if their pairwise distance exceeds the ANI threshold for the current hierarchical level. This is achieved by querying the edges table with dynamic SQL, filtering for node pairs present in the current reference set and whose distance satisfies the level threshold.

Once the graph is constructed, BacTaxID searches for maximal cliques, subgraphs where every node is connected to every other node, using the *maximal_cliques* algorithm from the petgraph library. A clique represents a set of genomes that are all mutually similar above the specified distance threshold, ensuring high internal consistency and preventing the chaining artifacts common in single-linkage or agglomerative clustering methods.

The algorithm iterates through all identified maximal cliques and evaluates them against the *click_size* parameter (typically 5 genomes). Because the classification process is dynamic and genomes are added sequentially, it is highly unusual for more than one new clique to form simultaneously. The introduction of a single new query genome is typically insufficient to bridge two previously separate, unclassified groups into distinct, valid cliques at the same time. Therefore, the algorithm considers the first clique found that meets or exceeds the minimum size as the valid new cluster. This “first-found” heuristic balances computational cost with biological representativeness, as larger cliques typically represent well-established, densely populated regions of the genomic landscape, and the sequential nature of the algorithm makes the discovery of multiple concurrent cliques a rare event.

Upon identification of a valid clique, a new hierarchical code is generated for that level. The system queries the code table to determine the next available integer code for the current parent group (e.g., if the highest existing code for parent 1.3 is 1.3.5, the new clique is assigned code 1.3.6). This new code is assigned to the query genome, and its status is set to classifier (C).

Crucially, this new classification is then propagated to all other members of the identified clique. Each genome within the clique has its hierarchical code updated in the database to reflect membership in the new cluster. Other genomes present in the initial distance graph but not part of the validated clique are also assigned the new code, but with a satellite (S) status. This ensures they are associated with the newly formed group but do not act as references in future classification steps.

To optimize subsequent hierarchical searches, the algorithm performs edge pruning. After a clique is processed at level j, all edges that constituted the clique are deleted from the edges table if their distance is below the threshold for the next finer level (j+1). This prevents redundant re-evaluation of these high-similarity relationships in subsequent, more stringent clique searches. This strategic pruning ensures that the edges table remains lean and that clique detection at finer resolutions operates on a reduced, relevant subset of the data, further enhancing computational scalability.

### Creation of the All the Bacteria Classification Scheme

BacTaxID classification schemes were generated for 67 bacterial and archaeal genera selected from the All the Bacteria (ATB) database^16^ based on the availability of ≥500 quality-controlled genomes per genus. The ATB collection comprised 2.4 million genome assemblies as of August 2024, yielding a total of 2.3 million genomes distributed across 67 genera and 3,926 species for hierarchical classification. Individual typing schemes were constructed for each genus using uniform parameterization (*sketch_size* = 3000, *kmer_size* = 31, *click_size* = 5, *click_threshold* = 0.8, *reference_size* = 100), with six hierarchical clustering levels corresponding to average nucleotide identity (ANI) thresholds of 96%, 98%, 99%, 99.5%, 99.9%, and 99.99%. Each genome was assigned a hierarchical typing code composed of up to six integers separated by periods (for example, 1.3.1.8.12.1), where each integer represents the cluster assignment at successive levels of resolution. All pre-computed schemes were deposited in a public repository and are freely accessible with accompanying metadata, classification results, and interactive exploration tools at www.bactaxid.org and https://zenodo.org/records/17Z917Z2.

### Statistical Analysis

Correlation between genome number within each genus and level-specific classification completeness was assessed using pearson correlation coefficient. Completeness was defined as the proportion of genomes assigned definitive hierarchical codes at each successive level of resolution.

Biological validity of the hierarchical framework was systematically evaluated by computing Normalized Mutual Information (NMI)^24^ between BacTaxID classifications at each hierarchical level and independent reference typing schemes, including MLST, core-genome MLST (cgMLST), and species designations. Initial validation analyses were performed on two model organisms, *Escherichia* and *Salmonella*, and subsequently extended across 11 representative genera (*Acinetobacter, Enterobacter, Enterococcus, Escherichia, Haemophilus, Klebsiella, Neisseria, Pseudomonas, Salmonella, Staphylococcus,* and *Streptococcus*), encompassing 1.7 million genomes representing approximately 74% of the All the Bacteria collection. NMI values were computed to quantify the degree of concordance between classification systems, with values ranging from 0 (independent) to 1 (perfect agreement), enabling direct comparison of clustering quality across hierarchical levels.

To further assess the discriminatory power and epidemiological relevance of BacTaxID classifications, a prevalence study simulation was conducted using *Escherichia coli* as a representative model organism. A random subset of 30,000 whole-genome sequences of E. coli was selected from the All the Bacteria collection and subjected to BacTaxID classification across all hierarchical levels.

To assess epidemiological utility, comprehensive outbreak detection analyses were performed on well-characterized nosocomial and zoonotic *Escherichia coli* transmission events with documented epidemiological linkage and independent molecular confirmation. For each outbreak, BacTaxID cluster assignments at all hierarchical levels were compared against published clinical and epidemiological definitions derived from core-genome MLST (cgMLST) profiles and pairwise single nucleotide polymorphism (SNP) distance matrices. Phylogenetic relationships among outbreak-associated isolates were inferred using UPGMA (Unweighted Pair Group Method with Arithmetic Mean)^50^ hierarchical clustering based on BacTaxID k-mer distance metrics, providing visual assessment of clonal structure and reconstruction of probable transmission chains.

Within-cluster SNP density was quantified by calculating pairwise SNP differences among isolates assigned to identical BacTaxID codes at hierarchical levels L_4_ (99.5% ANI) and L_5_ (99.99% ANI) across eight representative genera (*Acinetobacter, Enterobacter, Escherichia, Haemophilus, Klebsiella, Neisseria, Pseudomonas,* and *Salmonella*). SNPs between genome pairs were identified using the *snps_pairwise* function from pato^51^, which performs individual pairwise genome alignments to identify single nucleotide polymorphisms and calculate SNP density normalized to genome alignment. Results were expressed as SNPs per megabase (SNPs/Mb) of whole genome alignment, enabling standardized comparison across genera with varying genome sizes and evolutionary rates.

### Statistical Software and Visualization

All descriptive statistical analyses were conducted using R (version 4.5)^52^ with the following complementary packages: *tidyverse*^53^ for data manipulation and visualization pipelines, *DBI*^54,55^ and *duckdb*^47^ for efficient database query and integration, *ggsci*^56^ for scientific color palettes, *patchwork*^57^ for multi-panel figure composition, *ggpubr*^58^ for publication-ready graphical outputs, *tidyinftheo*^59,60^ for information-theoretic calculations, *ggrepel*^61^ for optimized text label placement, *ggnewscale*^62^ for multiple color scales within single plots, and *ggtree*^63–65^ for phylogenetic tree visualization and annotation.

Additionally, comprehensive interactive hierarchical classification visualizations were generated for all 68 bacterial and archaeal genera using KronaTools^66^, a specialized bioinformatics tool designed for multilevel taxonomic and metagenomic data visualization (available at https://github.com/marbl/Krona). KronaPlots enable rapid exploration of hierarchical BacTaxID classifications and provide intuitive assessment of genus-level and strain-level diversity across the complete taxonomy. All genus-specific KronaPlots have been integrated and are publicly accessible through the BacTaxID web platform at www.bactaxid.org, facilitating interactive data exploration and dynamic visualization of hierarchical classification schemes for researchers and public health practitioners.

### Genus-Specific Supplementary Typing Annotations

To provide enhanced epidemiological and evolutionary context, standardized typing annotations were obtained for select genera using specialized in silico typing resources:

#### Salmonella

Supplementary serovar, serotype, cgMLST and MLST were obtained through SISTR^27^ (*Salmonella* In Silico Typing Resource), which performs rapid in silico serotyping and subspecies assignment based on genomic sequences.

#### Klebsiella and Neisseria

Annotation data including strain typing and virulence factors were sourced from the All the Bacteria supplementary resource, available at https://allthebacteria.readthedocs.io/en/latest/typing.html.

#### Escherichia

Phylogroups were determined in silico using ClermontTyping^67^, a specialized classification tool designed for rapid phylogenetic assignment of *Escherichia* isolates into species and recognized phylogroups (A, B1, B2, D, E, F, and cryptic), accessible at https://github.com/A-BN/ClermonTyping.

Multilocus Sequence Typing (MLST) for bacterial genera not subjected to specialized typing analysis, conventional MLST sequence typing was performed using the standardized pipeline available at https://github.com/tseemann/mlst. This tool performs allelic profiling against appropriate species-specific MLST schemes and assigns sequence types (STs) according to established PubMLST^68,69^ nomenclature.

### cgMLST Genome Selection and Dataset Preparation

To evaluate the correlation between BacTaxID taxonomic levels and cgMLST genetic distances, we created a homogeneous dataset of whole-genome sequences for each of the fourteen selected bacterial genera. Genome selection was performed through a stratified sampling approach to ensure comprehensive representation across all BacTaxID hierarchical levels. First, we filtered the complete dataset to include only genomes belonging to the largest BacTaxID L_0_ group (1.x.x.x.x.x), which contains the major bacterial species of interest. Subsequently, we employed a multi-level sampling strategy to select genomes across each BacTaxID level (L_1_ through L_5_). For each BacTaxID taxonomic level, we randomly selected n=10 distinct groups and sampled n=50 genomes per group, ensuring balanced representation across the hierarchical structure. This approach was implemented using a custom R function (*sample_levels*) that iteratively sampled genomes at each BacTaxID level, filtering for non-empty entries and avoiding duplicate selections.

### cgMLST Scheme Development and Allele Calling

Homemade cgMLST schemes were created for all 14 bacterial genera using ChewBBACA (v3.5.0)^70,71^ a widely-used software tool for schema creation and allele calling in bacterial genomics. Genome assemblies (in FASTA format) were input into ChewBBACA following standard quality control procedures. The software performed allele calling by comparing each genome against the genus-specific core gene set, assigning allelic profiles based on nucleotide variations in the loci. cgMLST was defined as the set of genes present in at least 90% of the genomes. The resulting cgMLST schemes exhibited variable sizes across genera, ranging from 714 genes (*Enterococcus faecium*) to 1,291 genes (*Campylobacter jejuni*), reflecting the genomic diversity and core genome composition of each bacterial taxon. All allele calling results were retained for downstream analysis, including isolates with incomplete allelic assignments.

### Comparative Analysis of BacTaxID Levels and cgMLST Distances

For each bacterial genus, we calculated the percentage of common alleles between all pairwise isolate comparisons at each BacTaxID level (L_1_ to L_5_) using hamming distance. Common alleles were defined as identical allelic assignments at homologous loci. We stratified the analysis by BacTaxID level to assess how taxonomic granularity correlates with genetic distance as measured by cgMLST. Isolates assigned to the same BacTaxID L_5_ group (the most specific BacTaxID level) were compared to quantify the number of differing alleles, providing a genus-specific resolution metric. Results were visualized using box plots to display the distribution of values (median, interquartile range, and extremes) across taxonomic levels and bacterial genera.

### Software availability and web resources

Pre-computed hierarchical classification schemes for all 68 bacterial and archaeal genera are publicly accessible without restrictions through the BacTaxID web platform (www.bactaxid.org), with full classification data, metadata, and interactive visualization tools provided at no cost. Complete datasets, including genomic identifiers, hierarchical codes, and supplementary annotations for all 2.3 million genomes in the All the Bacteria collection, have been deposited in the Zenodo research repository (https://zenodo.org/records/17791772) to ensure long-term preservation and accessibility.

The BacTaxID software framework, including source code and precompiled binaries, is available through open-source distribution at https://github.com/irycisBioinfo/BacTaxID under the GPL-3.0 license. Reproducible analysis scripts, complete data processing workflows, statistical analyses, and supplementary metadata tables supporting the results presented in this study are publicly available at https://zenodo.org/uploads/18493374.

## Acknowledgements and Funding

V.F.L. is supported by the “Miguel Servet program” (CP22/00164) from the Instituto de Salud Carlos III (ISCIII), designed to foster professional research careers within the Spanish National Health System (NHS).

M.D.F.d.B. was supported by the Instituto de Salud Carlos III (pFIS F19/00366), and V.F.L. by Miguel Servet funding (CP22/00164). This research received financial support from the European Commission (MISTAR AC21_2/00041), the Instituto de Salud Carlos III (ISCIII) through CIBERINFEC (CB21/13/00084), and the Fundación Francisco Soria Melguizo (CC23140547). During this study, M.D.F.d.B. was additionally supported by the pFIS program (F19/00366) from the ISCIII.

---

**Supplementary Figure 1.**
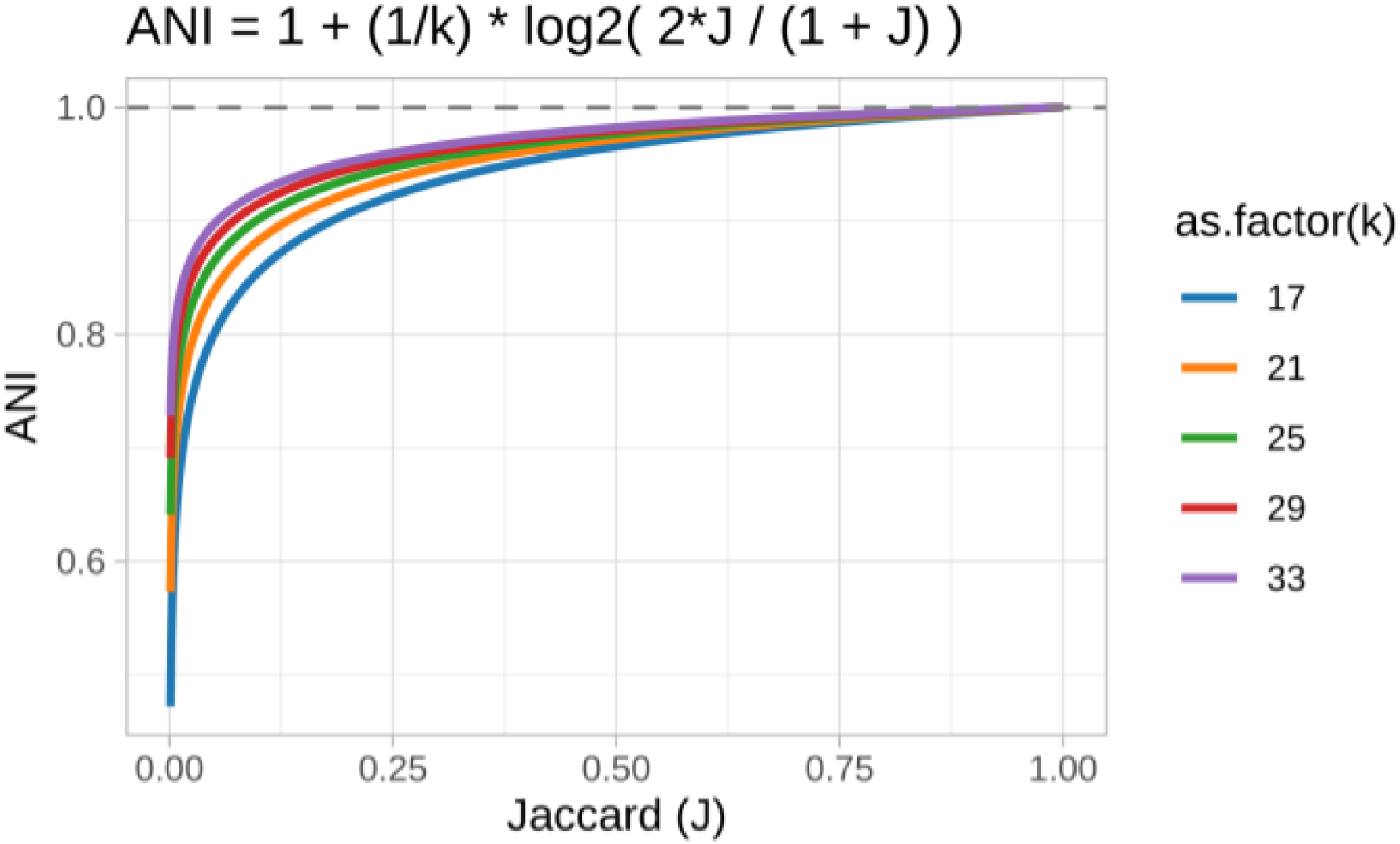
Theoretical relationship between the Jaccard index (J) and the Average Nucleotide Identity (ANI). The plot illustrates the Mash-derived equation for estimating ANI from J, a k-mer based similarity metric. Each colored line represents the estimated ANI value for a given Jaccard index, calculated using different k-mer sizes (k), ranging from 17 to 33. The dashed line indicates the theoretical maximum of 1.0 for both ANI and J.

